# Cigarette smoking and personality: Investigating causality using Mendelian randomization

**DOI:** 10.1101/246181

**Authors:** Hannah M Sallis, George Davey Smith, Marcus R Munafò

## Abstract

**Background:** Despite the well-documented association between smoking and personality traits such as neuroticism and extraversion, little is known about the potential causal nature of these findings. If it were possible to unpick the association between personality and smoking, it may be possible to develop more targeted smoking cessation programmes that could lead to both improved uptake and efficacy.

**Methods:** Recent genome-wide association studies (GWAS) have identified variants robustly associated with both smoking phenotypes and personality traits. Here we use publicly available GWAS summary statistics in addition to data from UK Biobank to investigate the link between smoking and personality. We first estimated genetic overlap between traits using LD score regression and then applied both one- and two-sample Mendelian randomization methods to unpick the nature of this relationship.

**Results:** We found clear evidence of a modest genetic correlation between smoking behaviours and both neuroticism and extraversion, suggesting shared genetic aetiology. We found some evidence to suggest an association between neuroticism and increased smoking initiation. We also found some evidence that personality traits appear to be causally linked to certain smoking phenotypes: higher neuroticism and heavier cigarette consumption, and higher extraversion and increased odds of smoking initiation. The latter finding could lead to more targeted smoking prevention programmes.

**Conclusion:** The association between neuroticism and cigarette consumption lends support to the self-medication hypothesis, while the association between extraversion and smoking initiation could lead to more targeted smoking prevention programmes.

## Introduction

There is a well-documented association between smoking and personality traits such as neuroticism and extraversion (Terracciano & Costa Jr. 2004; Malouff et al. 2006; Munafò et al. 2007; Hakulinen et al. 2015), and with associated mental health outcomes such as major depressive disorder (MDD) (Munafò & Araya 2010; Fluharty *et al.* 2017). However, given that much of these data come from observational studies, it is difficult to make any causal inference regarding these relationships. It is possible that the observed associations could be due to confounding, and if a true causal relationship does exist the direction of effect is unknown.

Although we now know that smoking has a causal effect on mortality, there has previously been some discussion surrounding this. It had been suggested by Eysenck and others that personality influences both mortality and smoking independently, and that this leads to a non-causal association between smoking and mortality (Eysenck 1965). Much work went into investigating the Type A personality type, which was argued to be a risk factor for coronary heart disease and other health outcomes, although many of these findings failed to replicate (Petticrew et al. 2012).

Understanding these relationships is therefore important for public health and policy. The World Health Organisation (WHO) now recognises smoking as one of the leading modifiable risk factors for disability, disease and death (World Health Organisation 2002). As a result, if it were possible to unpick the association between personality and smoking, it may be possible to develop more targeted smoking cessation programmes, which could lead to both improved uptake and efficacy. It has also been suggested that personality traits are potential modifiable targets for intervention. Understanding the nature of these associations could therefore lead to novel interventions addressing the specific traits associated with smoking behaviours (Roberts & Hill 2017).

Neuroticism and extraversion are two of the main components of personality. The former reflects emotional instability, stress-vulnerability and proneness to anxiety (Kendler et al. 1993). Higher neuroticism has been linked to anxiety and MDD, with some evidence of shared genetics and a causal link between neuroticism and MDD onset (Neale et al. 2005; Gale et al. 2016). Although levels of neuroticism are increased among smokers, the evidence that neuroticism is linked with smoking initiation is inconsistent, with one meta-analysis suggesting that neuroticism is linked with relapse to smoking among former smokers (Hakulinen et al. 2015) rather than smoking initiation. In contrast, extraversion is characterised by tendencies such as liveliness and assertiveness of an individual and the level of ease and enjoyment of social interactions (Kendler et al. 1993; van den Berg et al. 2016). There is some suggestion in the literature that high levels of extraversion are associated with greater rates of smoking initiation, and lower rates of smoking cessation (Hakulinen et al. 2015).

Mendelian randomization (MR) is a method of assessing causality from observational data through the use of genetic instrumental variables (Sallis et al. 2014). Genetic variants (or scores constructed from several variants) are used as a proxy for some modifiable risk factor, for example smoking (Figure 1). The MR principle relies on an approximation of Mendel’s first and second laws:, that genotypes transmit across conception to a viable conceptus, independent of both environment and other genetic variants (Davey Smith 2011). Assuming the genetic variants are not associated with the outcome other than through the risk factor they act as a proxy for, we can make inferences about the causal direction of any association between the risk factor and the outcome (Davey Smith & Ebrahim 2003). If the underlying assumptions of MR are satisfied, the resulting effects given by the MR analysis should be free from the problems of confounding and reverse causality to which observational epidemiology is prone (Sallis et al. 2014).

**Figure 1.**
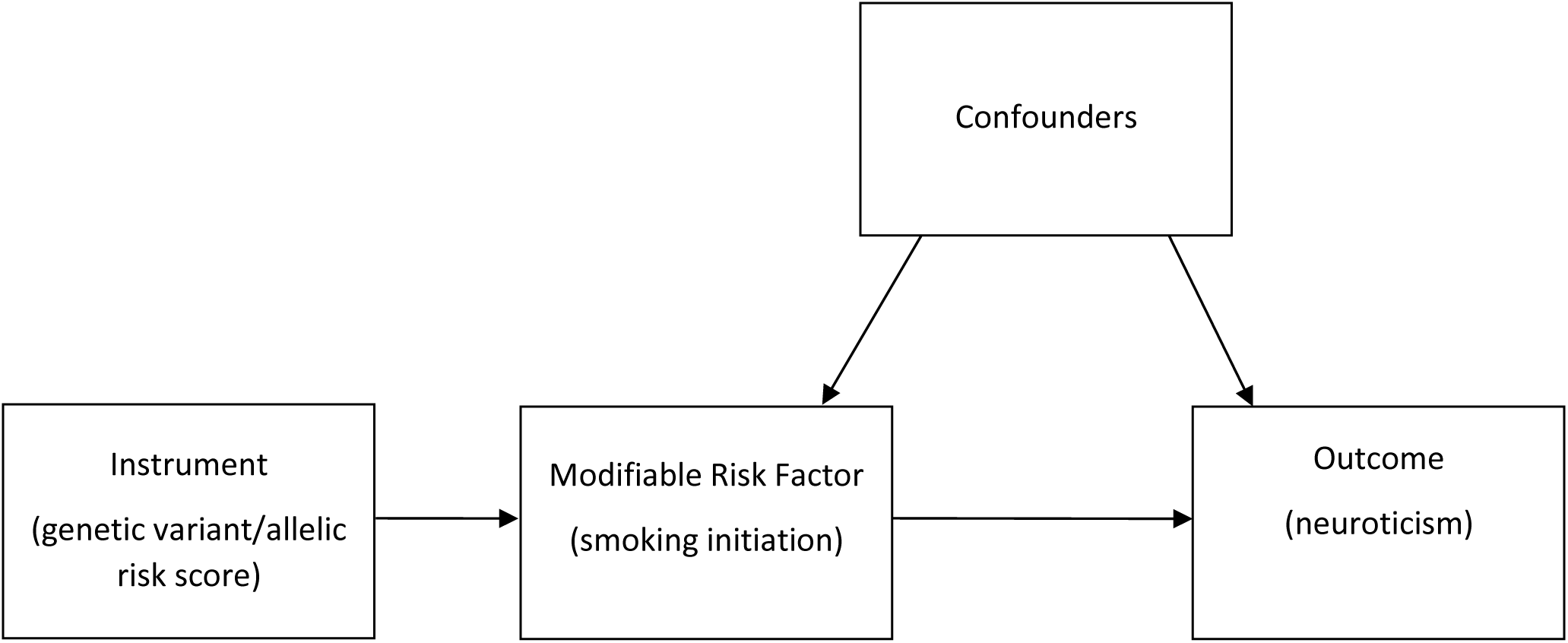
Directed acyclic graph illustrating Mendelian randomization. In this model, allelic risk scores associated with smoking initiation are calculated and used to assess the association of smoking initiation with neuroticism levels.

Previous work using MR found no evidence of a causal relationship in the direction of smoking to depression (Taylor et al. 2014). We therefore hypothesised that if a causal association with neuroticism exists, it is more likely to act from neuroticism to smoking. Methods have also been developed to investigate genetic correlation between traits. Although these do not provide information on causality of any potential relationship, they shed light on the amount of shared genetic architecture across traits. Any overlap here could be due to pleiotropy (genetic effects on multiple traits), shared biological mechanisms between traits, or a causal relationship from one trait to another but the direction of this cannot be ascertained from these approaches. Pleiotropy can be either biological, with variants associated with multiple independent traits, or mediated, where variants are associated with multiple phenotypes on the same causal pathway (Solovieff et al. 2013). Recent genome-wide association studies (GWAS) have identified variants robustly associated with a number of smoking phenotypes (The Tobacco and Genetics Consortium 2010) and with personality traits (Okbay et al. 2016; van den Berg et al. 2016). The availability of these summary statistics enables us to look at the extent of genetic correlation between smoking phenotypes and personality. This can be followed up by a range of MR methods to try and to unpick the nature of this relationship. Here we use publicly available GWAS summary statistics in addition to data from UK Biobank (UKB) (Sudlow et al. 2015) in a bidirectional analysis to investigate whether there appears to be a causal link between smoking and personality.

## Methods and measures

MR techniques using both genome-wide summary statistics and individual level data were used to investigate whether observed associations between smoking and personality traits are causal.

### Genetic instruments

#### Smoking behaviours

For each of the smoking phenotypes, a single variant was used as a genetic instrument for this behaviour. These were the rs6265 variant in the *BDNF* gene identified for smoking initiation, the rs16969968 variant in the *CHRNA5* gene for smoking heaviness among past and current smokers, and the rs3025343 variant in the *DBH* gene for smoking cessation. For each phenotype, the effect allele was that which corresponded to an increase in the relevant smoking behaviour.

#### Personality

Eleven independent variants associated with neuroticism were reported by Okbay et al. (2016) Of the original variants, 5 of these were unavailable in the TAG smoking data. Proxies were identified for 4 of these SNPs using SNIPA (Arnold et al. 2015) (r^2^>0.85). A complete list of variants used in the neuroticism instrument can be found in Table S1. Five independent variants associated with extraversion were identified by the GPC (van den Berg et al. 2016). Of these variants, 3 variants were unavailable in the TAG summary statistics. Using SNIPA, we identified a proxy for one of these variants (Table S1).

### Data sources

#### GWAS summary statistics

Publicly available summary statistics are available for recent genome-wide analyses of both smoking phenotypes and personality traits. The Tobacco and Genetics (TAG) consortium performed GWAS on several measures related to cigarette smoking, including smoking initiation, cessation and heaviness (The Tobacco and Genetics Consortium 2010). The consortium identified several hits, including rs6265 in the *BDNF* gene for smoking initiation, rs16969968 in the *CHRNA5* gene for smoking heaviness and rs3025343 in the *DBH* gene for smoking cessation. A recent GWAS investigating subjective well-being, depression and neuroticism identified 11 independent variants associated with neuroticism (Okbay et al. 2016), while 5 variants associated with extraversion were reported by the Genetic of Personality Consortium (GPC) (van den Berg et al. 2016).

#### UK Biobank (UKB)

UKB has collected phenotypic information on around 500,000 participants, with genotyping available on approximately 337,106 unrelated Europeans, exclusion criteria and quality control measures are described in detail elsewhere (Bycroft et al. 2017; Mitchell et al. 2017). An interim release of genetic data was made available in 2015 for a subset of the cohort. This subset was included in the neuroticism GWAS and contained approximately 114,780 European individuals (Sudlow et al. 2015).

Smoking status was defined as ever (consisting of current and former smokers) or never smoker according to responses given at the initial assessment visit in UKB. Smoking heaviness was derived for former and current smokers based on responses to ‘number of cigarettes currently smoked daily’ at the initial assessment. For former smokers, this question related to number of cigarettes previously smoked daily.

Neuroticism scores were derived from a number of neurotic behaviour domains measured at the initial assessment visit. These scores were externally derived by Smith et al. (2013) and are available for use by researchers accessing the UKB resource. Scores range from 0 to 12 with a higher score corresponding to a greater number of neurotic behaviours. There was no direct measure of extraversion in UKB, so analyses of this trait were restricted to those based on the genetic instrument for extraversion.

Polygenic risk scores for neuroticism and extraversion were calculated for each individual in UKB. The neuroticism risk score ranged from 1 to 19 and corresponded to the number of neuroticism increasing alleles per individual. A risk score was calculated for extraversion and ranged from 0 to 6. Although weighted scores can give a more precise effect estimate, the neuroticism GWAS included the interim release of UKB within the discovery sample (Okbay et al. 2016; Major Depressive Disorder Working Group of the PGC et al. 2017). Risk scores should use weightings derived from independent samples to avoid introducing bias into the effect estimates (Hartwig & Davies 2016). As a sensitivity analysis, we restricted analyses using the neuroticism risk score to participants who were not included in the interim release of the genetic data, and who were therefore not included in the discovery sample.

### Statistical analysis

#### Genetic correlation

In a first step, GWAS summary statistics were used to estimate the genetic correlation of smoking initiation with both neuroticism and extraversion. LD score regression was performed (without constraining the intercept) using the GWAS summary statistics to assess the amount of genetic overlap between the two traits. In order to estimate genetic correlation between personality measures and additional smoking phenotypes of smoking heaviness and cessation, summary statistics for the personality GWAS would need to be stratified by smoking status. Although the original GWAS summary statistics were not available stratified by smoking status, it was possible to estimate these genetic correlations using the individual level data available in UKB. Genome-wide complex trait analysis (GCTA) software (Yang et al. 2011) was used to estimate genetic correlation for the additional smoking phenotypes within UKB.

#### Two-sample MR using summary statistics

Bidirectional two-sample MR analyses were performed using the genetic instruments described above. Effect estimates and standard errors (SEs) were extracted for each variant from the relevant GWAS results and used to estimate inverse variance weighted (IVW) effect estimates. For the neuroticism instrument which incorporated multiple SNPs, we performed sensitivity analyses. Effect estimates and SEs were extracted from the original GWAS results as described above and MR-Egger (Bowden et al. 2015) and weighted median regression (Bowden et al. 2016) approaches were used to calculate effect estimates adjusted for pleiotropy and invalid instruments. Neuroticism and extraversion GWAS results were not stratified by smoking status. As a result, when using summary statistics, analyses in the direction of smoking to personality were restricted to smoking initiation only.

#### One-sample MR using individual level data

Further analyses were performed using data from the UKB. In these analyses, the association between the genetic instrument (G) and the outcome (Y) was estimated (Y~G). Within UKB it was possible to stratify participants according to smoking status. Therefore, in addition to smoking initiation, we also investigated the association between both smoking heaviness and cessation with personality. These analyses were adjusted for the top 10 principal components as well as genotype array.

## Results

### Genetic correlation

LD score regression using summary statistics from the TAG consortium GWAS of smoking initiation and the Okbay et al. neuroticism GWAS suggested evidence of a modest genetic correlation between the two traits (rG=0.124, SE=0.05). There was also evidence of a larger genetic correlation between extraversion and smoking initiation (rG=0.288, SE=0.01; Table 1). We used GCTA software to calculate genetic correlation using individual level data from UKB. There was evidence of genetic correlation between neuroticism and smoking heaviness among both current (rG=0.248, SE=0.12) and former smokers (rG=0.220, SE=0.06). We found evidence of a negative genetic correlation between smoking cessation and neuroticism (rG=-0.314, SE=0.15; Table 1).

**Table 1.**
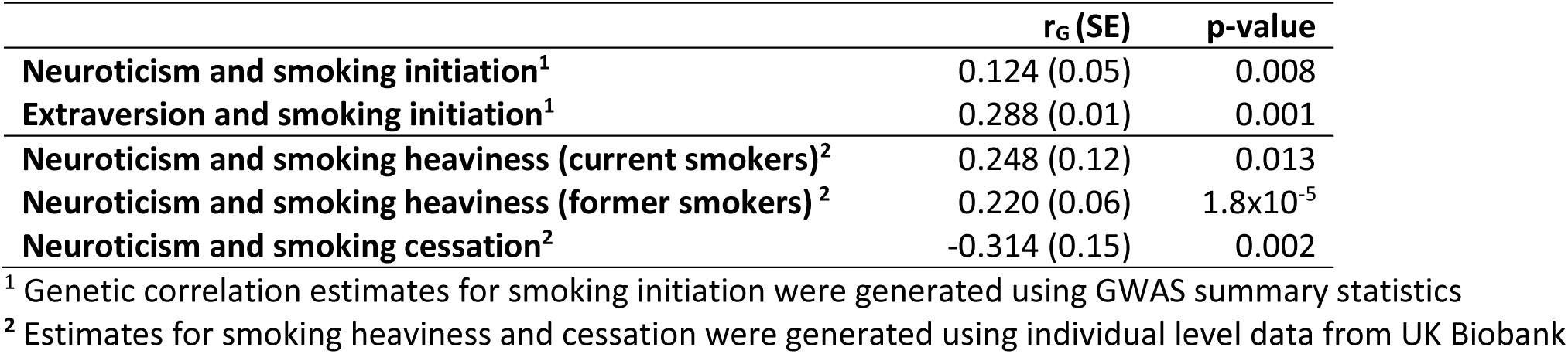
Genetic correlation between smoking phenotypes and personality traits using GWAS summary statistics and individual level data from UK Biobank.

### Observational association between neuroticism and smoking behaviours

Data on smoking status and neuroticism were available from UKB. A total of 389,770 participants had data available on both smoking and neuroticism, with genotyping available on 273,516 of these after applying QC measures. There was strong evidence of an observational association between neuroticism and smoking status in both the entire UKB sample and when restricting to those with genotyping data. Mean neuroticism scores were higher among former (4.22, SD=3.3) and current smokers (4.66, SD=3.5) than non-smokers (3.89, SD=3.2, p<0.001). We found evidence of an association between neuroticism and cigarettes smoked per day with heavier smokers reporting greater levels of neuroticism (β=0.02, p<0.001).

### Effects of smoking on personality traits

We first used two-sample MR to investigate the effect of smoking initiation on personality using publicly available GWAS summary statistics. This found no clear evidence of a relationship from smoking initiation to neuroticism when using rs6265 as an instrument for smoking initiation (β =-0.032, 95% CI: -0.16, 0.09, p=0.617; Table 2). A one-sample approach was also used to investigate the association between each smoking behaviour and neuroticism in UKB. This found weak evidence of association, with each copy of the smoking initiation risk allele associated with a decrease in neuroticism score (β=-0.023, 95% Cl: - 0.045, -0.001, p=0.037; Table 2).

**Table 2.** Effect of smoking on personality traits using one- and two-sample MR.

| Exposure | Outcome | Sample | Instrument | Smoking status | $\beta$ | 95% CI | P-value | N |
| --- | --- | --- | --- | --- | --- | --- | --- | --- |
| Smoking initiation | Neuroticism | GWAS summary statistics | rs6265 | All | -0.032 | -0.16, 0.09 | 0.617 | 170,911 |
|  |  | UK Biobank |  | All | -0.023 | -0.045, -0.001 | 0.037 | 274,230 |
|  | Extraversion | GWAS summary statistics | rs6265 | All | 0.198 | -0.03, 0.42 | 0.084 | 63,030 |
|  |  | UK Biobank |  |  | - | - | - | - |
| Smoking heaviness | Neuroticism | UK Biobank |  | Non-smokers | 0.004 | -0.02, 0.03 | 0.745 | 149,960 |
|  |  |  |  | Former smokers | 0.011 | -0.02, 0.04 | 0.472 | 96,674 |
|  |  |  |  | Current smokers | 0.038 | -0.02, 0.10 | 0.234 | 26,882 |
| Smoking cessation | Neuroticism | UK Biobank |  | Non-smokers | 0.004 | -0.03, 0.04 | 0.822 | 149,960 |
|  |  |  |  | Ever smokers | 0.029 | -0.01, 0.07 | 0.161 | 123,556 |

The genetic variant rs16969968 was used as a proxy for smoking heaviness in UKB. Despite strong evidence of an observational association between smoking heaviness and neuroticism, we found no robust evidence of a causal association from the rs16969968 variant for smoking heaviness to neuroticism among either former or current smokers (Table 2). Using the rs3025343 variant as a proxy for smoking cessation found no strong evidence of an association from smoking cessation to neuroticism (β=0.029, 95% CI: -0.01, 0.07, p=0.161; Table 2) in UKB.

When looking at the association between extraversion and smoking, we found weak evidence of an association between smoking initiation and increased extraversion (β=0.198, 95% Cl: -0.03, 0.42; Table 2) using two-sample MR. There was no relevant measure of extraversion in UKB, so we were unable to look at the association from smoking initiation to extraversion using individual level data in UKB.

### Effects of personality traits on smoking

Two-sample MR using summary statistics found no clear evidence of an association from neuroticism to smoking initiation (OR=1.165, 95% CI: 0.71, 1.91, p=0.499; Table 3). Several MR approaches were used to investigate the association from neuroticism to smoking initiation. MR-Egger and weighted median approaches found no robust evidence of an association after adjusting for pleiotropy and allowing for invalid instruments (Table S2). Among the UKB participants, there was no clear evidence of an association from neuroticism to smoking initiation when using an unweighted risk score (OR=1.000, 95% CI: 0.997, 1.003, p=0.980; Table 3). When looking at the association from extraversion to smoking initiation, we found no strong evidence of an effect when using a two-sample approach (OR=1.733, 95% Cl: 0.37, 8.23, p=0.268). However, this may be due to a lack of power. The direction of effect was consistent within the UKB sample, where there was strong evidence of an association. Each additional extraversion allele was associated with an increase in the odds of being an ever smoker (OR=1.015, 95% CI: 1.01, 1.02, p=9.6x10^-7^; Table 3).

**Table 3.** Effect of personality traits on smoking using one- and two-sample MR.

| Exposure | Outcome | Sample | Instrument | Smoking status | OR | 95% CI | P-value | N |
| --- | --- | --- | --- | --- | --- | --- | --- | --- |
| Neuroticism | Smoking initiation | GWAS summary statistics | IVW | All | 1.165 | 0.71, 1.91 | 0.499 | 74,035 |
|  |  | UK Biobank | Unweighted score | All | 1.000 | 0.997, 1.003 | 0.980 | 318,985 |
| Extraversion | Smoking initiation | GWAS summary statistics | IVW | All | 1.733 | 0.37, 8.23 | 0.268 | 74,035 |
|  |  | UK Biobank | Unweighted score | All | 1.015 | 1.01, 1.02 | 9.6x10<sup>-7</sup> | 332,596 |
|  |  |  |  |  | β | 95% CI | P-value | N |
| Neuroticism | Smoking heaviness | GWAS summary statistics | IVW | Ever smokers | 0.050 | -4.11, 4.21 | 0.979 | 38,181 |
|  |  | UK Biobank | Unweighted score | Former smokers | -0.018 | -0.05, 0.02 | 0.315 | 70,909 |
|  |  |  |  | Current smokers | 0.068 | 0.02, 0.12 | 0.009 | 21,449 |
| Extraversion | Smoking heaviness | GWAS summary statistics | IVW | Ever smokers | 0.017 | -16.33, 16.37 | 0.997 | 38,181 |
|  |  | UK Biobank | Unweighted score | Former smokers | -0.021 | -0.08, 0.04 | 0.519 | 73,967 |
|  |  |  |  | Current smokers | -0.038 | -0.13, 0.05 | 0.419 | 22,340 |
|  |  |  |  |  | OR | 95% CI | P-value | N |
| Neuroticism | Smoking cessation | GWAS summary statistics | IVW | Ever smokers | 0.797 | 0.43, 1.47 | 0.423 | 41,278 |
|  |  | UK Biobank | Unweighted score | Ever smokers | 0.997 | 0.99, 1.00 | 0.272 | 144,119 |
| Extraversion | Smoking cessation | GWAS summary statistics | IVW | Ever smokers | 0.586 | 0.07, 4.76 | 0.387 | 41,278 |
|  |  | UK Biobank | Unweighted score | Ever smokers | 0.989 | 0.98, 1.00 | 0.057 | 150,294 |

Using an IVW approach, we found no clear evidence of an association from neuroticism to smoking heaviness (β=0.050, 95% CI: -4.11, 4.21, p=0.979; Table 3). However, MR-Egger suggested some evidence of biological pleiotropy (β=-0.500, p=0.026; Table S2) and a bias adjusted estimate suggested some evidence of an association between neuroticism and increased smoking heaviness (β=22.55, p=0.027). We also looked at this association in UKB when stratifying according to smoking status and found some evidence of an association. Among current smokers, the neuroticism risk score was associated with increased smoking heaviness (β=0.068, 95% CI: 0.02, 0.12, p=0.009; Table 3). In this analysis, each additional neuroticism risk allele was associated with smoking an extra 0.07 cigarettes per day. We found no robust evidence of an association from extraversion to smoking heaviness when using a two-sample MR approach (β =0.017, 95% Cl: -16.33, 16.37, p=0.997; Table 3) or when stratifying on smoking status and investigating this association in UKB. This remained the case among both former (β=-0.021, 95% Cl: - 0.08, 0.04, p=0.519) and current smokers (β=-0.038, 95% CI: -0.13, 0.05, p=0.419; Table 3).

When using two-sample MR with summary statistics, we found no robust evidence of association between neuroticism and smoking cessation when using the IVW approach, or when adjusting for bias (Tables 3, S2). This remained the case when looking within UKB (OR=0.997, 95% CI: 0.99, 1.00, p=0.272; Table 3). There was no strong evidence of an effect from extraversion to smoking cessation when using two-sample MR with summary statistics (OR=0.586, 95% CI: 0.07, 4.76, p=0.387; Table 3). When restricting our analyses to current and former smokers within UKB, we found weak evidence of an association. Each additional increase in extraversion risk allele was associated with 1.1% lower odds of smoking cessation (p=0.057; Table 3).

### Sensitivity analyses

Analyses involving neuroticism were also performed restricting to participants whose genetic data was not included in the interim release of UKB data. Full results are reported in Table S3. Results remained largely consistent. In this subset, the strength of evidence for the effect of neuroticism on smoking heaviness was weakened (current smokers: β=0.053, p=0.120). However, the effect size remained consistent, so this may be due to a lack of power in this smaller sample.

## Discussion

We attempted to disentangle the relationship between smoking and the personality traits of neuroticism and extraversion. Although much of the observed association between smoking and personality appears to be non-causal, we found evidence of a modest genetic correlation with both neuroticism and extraversion, suggesting some shared genetic aetiology. Given that available GWAS summary statistics for neuroticism and extraversion are not stratified by smoking status, we initially used two-sample MR approaches to look at the bidirectional association with smoking initiation. This was followed by one-sample MR using individual level data from UKB. When looking at the association from smoking to personality, we found some evidence that the rs6265 variant for smoking initiation was associated with both decreased neuroticism and increased extraversion. When looking in the other direction, we found evidence that the neuroticism risk score was associated with increased smoking heaviness, and that the extraversion risk score was associated with increased smoking initiation and decreased smoking cessation.

Both two-sample MR and one-sample analyses in UKB found consistent effects of the smoking initiation instrument (rs6265) on lowering neuroticism scores, although the evidence for this was weak. Although this is a strong instrument for smoking initiation, there could be potential pleiotropic effects. The inclusion of several strongly associated, but independent variants could reduce the potential impact of these effects, as all pleiotropic effects would need to operate in the same direction (Gage et al. 2017). Analyses from personality to smoking behaviours found some evidence of an association between increased genetic liability for neuroticism and greater smoking heaviness using one-sample MR and two-sample MR after adjusting for biological pleiotropy. Both the observational and MR analyses found a stronger effect among current smokers, where at least part of this association appears to be a causal effect. The observed association between smoking initiation and a decrease in neuroticism scores, plus the association between increased levels of neuroticism and heavier smoking would appear to lend support to the self-medication hypothesis.

We also observed evidence of an association between extraversion and smoking. Unlike neuroticism, extraversion did not show evidence of a causal relationship with smoking heaviness, but we did find an association with smoking initiation. Although there was no strong evidence of an association when using a two-sample approach, this could be due to a lack of power given that the direction of effect was consistent with that observed in UKB. Using UKB data, there was evidence that individuals with a higher genetic liability for extraversion had greater odds of taking up smoking. One potential mechanism for this is that extraversion could lead to more social contacts and greater susceptibility to peer influences, which are known to be important in smoking initiation. These findings could be taken forward to develop novel interventions. If self-medication does contribute to the smoking behaviours of individuals, as suggested by these results, it seems likely that targeting relevant personality traits, in addition to addressing the ensuing smoking behaviours could result in increased efficacy of any intervention.

There are a number of limitations to our analysis that should be considered. First, UKB formed a large part of the discovery cohort for the GWAS of neuroticism. We were therefore unable to use weighted risk scores to assess the association between smoking phenotypes and neuroticism in our one sample analyses – weights should be identified in independent samples to avoid overfitting the data and introducing bias into effect estimates (Hartwig & Davies 2016). However, we performed sensitivity analyses restricting to individuals who were not included in the discovery samples, and results remained consistent. Second, we were unable to use two-sample methods to assess the association from smoking heaviness and cessation to neuroticism and extraversion because the personality summary statistics were not stratified by smoking status. However, we did investigate the association in both directions for neuroticism when using the UKB data. Both the two-sample and UKB analyses gave consistent results when looking at the neuroticism to smoking initiation relationship. Third, due to the lack of an extraversion phenotype currently available in UKB we were unable to investigate whether there was evidence of an effect from smoking to extraversion. Fourth, MR analyses can often suffer from a lack of power, with large sample sizes and strong instruments required to detect effects. We have identified genetic variants robustly associated with each trait of interest based on results of large recently published GWAS in order to maximise the strength of our instruments. In this analysis, we use a combination of one- and two-sample MR approaches based on GWAS summary statistics in addition to data from UKB in order to maximise our power to detect any effect. Fifth, we stratified on smoking status to investigate the association of smoking heaviness and cessation phenotypes. Although this allows us to investigate pleiotropy, there is the potential to introduce collider bias (Munafò et al. 2017). However, our instruments are principally associated with smoking heaviness and cessation rather than smoking initiation, so that the risk of collider bias is minimised (Gage et al. 2016).

In conclusion, we found evidence of modest genetic correlation with both neuroticism and extraversion, suggesting some shared genetic aetiology and implying that much of the observed association between smoking and personality is non-causal. However, we also found some evidence for specific causal pathways between personality and smoking phenotypes - higher neuroticism and heavier cigarette consumption, and higher extraversion and increased odds of smoking initiation. The association between neuroticism and cigarette consumption lends support to the self-medication hypothesis, while the association between extraversion and smoking initiation could lead to more targeted smoking prevention programmes.

## Conflict of interest

None.

## Notes

Financial support: This work was supported by the Medical Research Council and the University of Bristol (MC_UU_12013/1, MC_UU_12013/6). MRM and HMS are members of the UK Centre for Tobacco and Alcohol Studies, a UKCRC Public Health Research: Centre of Excellence.

